# Antibodies against egg- and cell-grown influenza A(H3N2) viruses in adults hospitalized during the 2017-2018 season

**DOI:** 10.1101/439471

**Authors:** Min Z. Levine, Emily T. Martin, Joshua G. Petrie, Adam S. Lauring, Crystal Holiday, Stacie Jefferson, William J. Fitzsimmons, Emileigh Johnson, Jill M. Ferdinands, Arnold S. Monto

## Abstract

**Background:** The 2017-2018 US influenza season was severe with low vaccine effectiveness. Circulating A(H3N2) viruses from multiple genetic groups were antigenically similar to cell-grown vaccine strains. However, most influenza vaccines are egg-propagated.

**Methods:** Serum was collected shortly after illness onset from 15 PCR confirmed A(H3N2) infected cases and 15 uninfected (controls) hospitalized adults enrolled in an influenza vaccine effectiveness study.

Geometric mean titers against egg- and cell-grown A/Hong Kong/4801/2014 A(H3N2) vaccine strains and representative circulating viruses (including A/Washington/16/2017) were determined by microneutralization (MN) assays. Independent effects of strain-specific titers on susceptibility were estimated by logistic regression.

**Results:** MN titers against egg-A/Hong Kong were significantly higher among those who were vaccinated (MN GMT: 173 vs 41; *P* = 0.01). However, antibody titers to cell-grown viruses were much lower in all individuals (*P*>0.05) regardless of vaccination. In unadjusted models, a 2-fold increase in MN titers against egg-A/Hong Kong was not significantly protective against infection (29% reduction; p=0.09), but a similar increase in cell-A/Washington titer (3C.2a2) was protective (60% reduction; p=0.02). A similar increase in egg-A/Hong Kong titer was not significantly associated with odds of infection when adjusting for MN titers against A/Washington (15% reduction; P=0.61). A 54% reduction of odds of infection was observed with a 2-fold increase in A/Washington (not significant; P=0.07), adjusted for egg-A/Hong Kong titer.

**Conclusion:** Although individuals vaccinated in 2017-2018 had high antibody titers against the egg-adapted vaccine strain, antibody responses to cell-grown circulating viruses may not be sufficient to provide protection, likely due to egg-adaptation in the vaccine.

---

The 2017-2018 influenza season was severe in the United States, resulting in the highest rate of influenza-related hospitalizations since the Centers for Disease Control and Prevention (CDC) began including adults in inpatient surveillance[1]. Although the influenza A(H3N2) viruses that circulated in the US belonged to multiple genetic groups, they were all considered antigenically similar to cell-grown vaccine strain reference viruses[2]. However, the majority of available influenza vaccines are produced in eggs and vaccine effectiveness (VE) was low in preventing adult influenza A(H3N2) infections in 2017-2018 [3]. While the inadequate VE against A(H3N2) viruses was highlighted by the intensity of the 2017-2018 outbreak, the national level estimates for A(H3N2) vaccine effectiveness were consistent with observations of relatively low VE against this subtype throughout the past decade [4].

A common factor over the last several years of low VE against influenza A(H3N2) is that changes to the virus occur during its adaptation to eggs for vaccine production. Studies have suggested that changes required to adapt influenza viruses for growth in eggs may have contributed to low VE in years with influenza A(H3N2) circulation, including 2012-2013, 2016-2017, and this most recent season [5–7]. Egg-related changes can happen in multiple hemagglutinin epitopes and those egg-adaptation sites of concern have changed since the 2014-2015 season. Specifically, influenza A(H3N2) viruses belonging to the 3C.2a genetic group, which emerged in 2014-2015, acquired a glycosylation site in the hemagglutinin antigenic site B [6–8]. This glycosylation site is lost during adaptation to eggs, and, as a result, human and ferret serum raised against egg-adapted A(H3N2) vaccines have shown reduced inhibition of recently circulating viruses [7,9].

During the severe 2017-2018 influenza season, we carried out a study of influenza vaccine effectiveness among inpatients with acute respiratory illness (ARI) as part of the Centers for Disease Control and Prevention (CDC) sponsored Hospitalized Adult Influenza Vaccine Effectiveness Network (HAIVEN).

Preliminary vaccine effectiveness for the prevention of hospitalization due to influenza was estimated to be 16% and was not statistically significant [10]. At the HAIVEN site in Michigan, serum specimens were collected shortly after illness onset, representing pre-infection serologic immune status. Associations between antibody titers and influenza vaccination and A(H3N2) infection status were assessed, comparing antibody against egg- and cell-grown vaccine strains and a panel of wild-type viruses representing those that circulated.

## METHODS

### Study Population

Adult (≥18 years) patients hospitalized for treatment of acute respiratory illnesses at the University of Michigan (UM) Hospital in Ann Arbor were prospectively enrolled in a case-test negative design study of influenza vaccine effectiveness during the 2017-2018 influenza season using previously described methods [11]. All patients were enrolled ≤10 days from illness onset during active influenza circulation. Participants provided informed consent, completed an enrollment interview, and had throat and nasal swab specimens collected for influenza identification by RT-PCR using the CDC Influenza Division protocol. As part of the enrollment interview, participants rated their general health prior to the onset of their current illness (excellent, very good, good, fair, poor), and responded to a series of 5 questions assessing difficulties with various tasks to define frailty status[11,12]. General health was treated as a dichotomous variable (≥good vs. ≤fair), and participants were considered frail if they reported difficulties in ≥3 of the 5 elements of the frailty interview. Influenza vaccination status was documented by electronic medical record and state vaccination registry.

The institutional review board at the University of Michigan Medical School and Centers for Disease Control and Prevention reviewed and approved the study.

### Serologic analysis

Available residual clinical serum specimens collected after hospital admission were retrieved (0-9 days after illness onset). Specimens were considered to represent pre-infection serologic status, prior to a de novo antibody response to the incident illness. Among participants enrolled as of January 31, 2018 with available serum specimens, we selected all 15 influenza A(H3N2) positive cases identified to date and 15 influenza negative controls matched (nearest neighbor) on age and date of enrollment.

Microneutralization assays (MN) were performed using MDCK-SIAT1 cells [13,14] Briefly, sera were heat inactivated and twofold serial diluted, then mixed with 100 50% tissue culture infection dose (TCID50) of A(H3N2) viruses and incubated at 37°C 5% CO_2_ for 1 hr. The virus-sera mixture was used to infect 1.5 × 10^4^/well Madin-Darby Canine Kidney (MDCK)-SIAT1 cells and incubated for 18-20 hrs at 37°C with 5% CO_2_. After cold acetone fixation, the presence of viral protein was quantified by an ELISA using monoclonal antibodies specific to the nucleoproteins (NP) of the influenza A viruses. MN was conducted in the Influenza Division research laboratory at the Centers for Disease Control and Prevention. MN antibody titers were measured against influenza A(H3N2) viruses representing the egg and cell-culture grown A/Hong Kong/4801/2014 virus included in 2017-2018 Northern hemisphere influenza vaccines and 4 wild-type viruses representative of circulating influenza A(H3N2) genetic groups (Table 1).

**Table 1.** Influenza A(H3N2) virus strains used as targets in serologic assays

| Influenza A(H3N2) Strain | Propagation | Genetic Group | Percent of viruses identified in study surveillance <sup>b</sup> |
| --- | --- | --- | --- |
| A/HongKong/4801/2014 | Egg | 3C.2a | 0% |
| A/HongKong/4801/2014<sup>a</sup> | MDCK-SIAT1 | 3C.2a | 0% |
| A/Washington/16/2017 | MDCK-SIAT1 | 3C.2a2 | 83% |
| A/Singapore/Infimh-16-0019/2016 <sup>a</sup> | MDCK-SIAT1 | 3C.2a1 | 4% |
| A/Maryland/53/2017 | MDCK-SIAT1 | 3C.2a1b+135N | 0% |
| A/Wisconsin/327/2017 | MDCK-SIAT1 | 3C.2a1b+135K | 0% |

<sup>a</sup>NAI assay target
<sup>b</sup> 13% of sequenced viruses belonged to the 3C.3a genetic group, but all were isolated in April 2018, well after the influenza A(H3N2) cases included in serologic analyses were identified.

Neuraminidase-inhibition (NAI) antibody titers were determined for all 30 serum specimens collected from case and control participants using enzyme-linked lectin assays (ELLA) in the Respiratory Virus Laboratory at the University of Michigan School of Public Health[15]. NAI titers were measured using H6 reassortant influenza viruses (kindly provided by M. Eichelberger and H. Wan) bearing NA representing the A/Hong Kong/4801/2014 (H3N2) vaccine in 2017-18 season and A/Singapore/INFIMH-16-0019/2016 (H3N2) strains.

Antibody titers were calculated as the reciprocal of the highest dilution that neutralized 50% of virus infectivity (MN) or inhibited NA activity (ELLA). Titers below the limits of detection for each assay (i.e., <10) were denoted as half the threshold value (i.e., 5).

### Sequencing, phylogenetics and structure analysis

To characterize the viruses circulating in our source population during the study period, we identified all 29 A(H3N2) positive specimens with RT-PCR cycle threshold < 30 from University of Michigan participants enrolled during the full surveillance period (November 27, 2017, April 24, 2018). Six of these 29 were included as cases in the serologic analyses described above; the remaining 9 cases had RT-PCR cycle threshold ≥30. Segment 4 was amplified from extracted viral RNA using the SuperScript III One-Step RT-PCR Platinum Taq HiFi Kit (Invitrogen 12574) and primers HA_H3N2_F (5’ AGCAAAAGCAGGGGATAATTCTATTAACCATG 3’) and HA_H3N2_R (5’ AGTAGAAACAAGGGTGTTTTTAATTAATGCACTC 3’). The thermocycler protocol was: 50 °C for 60 min then 94 °C for 2 min then 30 cycles of 94 °C for 30 sec, 54 °C for 30 sec, 68 °C for 3 min, followed by a final 5 min extension at 68°C. All RT-PCR products were purified prior to Sanger sequencing. Twenty-three samples yielded sequence of sufficient quality for subsequent analysis.

Additional reference sequences representing circulating genetic groups were identified using publicly available sources [16] and downloaded from GISAID and GenBank (for all strains and reference numbers see Supplementary Table 1). Sequences were aligned using MUSCLE [17] with default parameters in MegAlign Pro Version: 13.0.0 (Lasergene). The alignments were trimmed to the HA coding region using Aliview v.1.23 [18]. Nucleotide substitution model selection was performed using jmodeltest v2.1.10 [19]. Maximum likelihood phylogenetic trees were generated in RAxML v8 [20] with a GTR gamma model and 1000 bootstraps. Trees were visualized and edited using FigTree v1.4.2

HA structure was generated using the PyMOL (Molecular Graphics System, Schrödinger, LLC)

### Statistical analysis

Individual antibody titers were log2 transformed for analysis; original values were divided by half the threshold value of detection (i.e. 5) to set the starting point of the log scale to zero prior to transformation. Mean log2 titers were calculated and compared by case status and influenza vaccination status using T-tests. The correlation of log2 titers against each virus were assessed using Spearman rank correlation coefficients (ρ). The independent effects of strain-specific MN titers on protection from infection was estimated in logistic regression models with RT-PCR-confirmed influenza A(H3N2) infection as the outcome and log2 MN titers as continuous predictors. Specific MN titers included in this model included those measured against the egg-grown A/Hong Kong/4801/2014 A(H3N2) vaccine strain and the A/Washington/16/2017 A(H3N2) circulating strain most closely related to the viruses that circulated in the local area. Odds ratios obtained from the model are interpreted as the factor reduction in the odds of influenza A(H3N2) infection associated with a 2-fold increase in MN titer against A/Hong Kong/4801/2014 A(H3N2) holding the A/Washington/16/2017 A(H3N2)-specific antibody titer constant, or vice versa.

Statistical analyses were carried out using SAS software (release 9.4; SAS Institute) and R (version 3.4.3; packages: ggplot2); a P value of <.05 or a positive lower bound of a 95% confidence interval (CI) were considered to indicate statistical significance.

## RESULTS

### Genetic characterization of circulating viruses

Influenza A(H3N2) viruses from 23 hospitalized cases with RT-PCR cycle threshold <30 were successfully sequenced to determine the characteristics of the strains circulating locally. Analyses determined that 19 viruses (83%) belonged to the 3C.2a2 genetic group similar to A/Washington/16/2017 (Figure 1). A single virus belonged to the 3C.2a1a group similar to A/Singapore/Infimh-16-0019/2016. Three viruses (13%) belonging to the 3C.3a genetic group were also identified, all of which were isolated in April 2018. Viruses descendent from the 3C.2a genetic group and serum collected during the first half of the influenza outbreak were used in serological analysis (Table 1) Compared to wild type SIAT1-grown virus A/Washington/16/2017 and SIAT1 grown-A/Hong Kong/4801/2014, vaccine-like egg propagated A/Hong Kong/4801/2014 has egg-adapted changes at T160K (antigenic site B, causing a loss of glycosylation motif), L194P (antigenic site B) and N96S (antigenic site D) (Figure 2).

**Figure 1.**
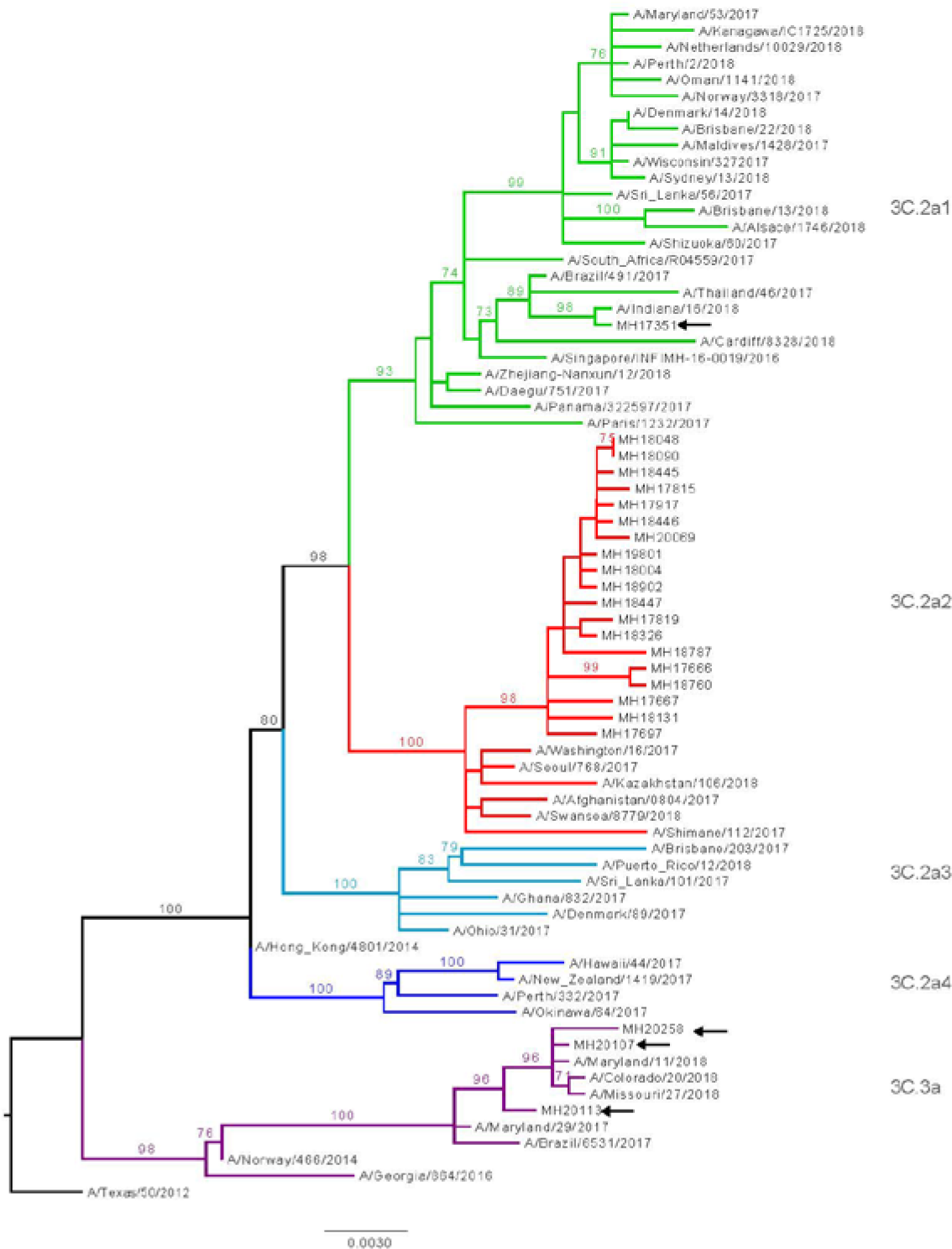
Maximum likelihood trees of HA sequences with branches color-coded by clade (3C.2a1-3C.2a4 and 3C.3a, see notation near tips). The outgroup is A/Texas/50/2012. Samples from the HAIVEN cohort begin with “MH,” see clade 3C.2a2 (n=19) and 4 additional sequences (Black Arrows). Clade assignments are based on locations of the reference sequences in nextstrain.org. Bootstrap values ≥ 70 are shown.

**Figure 2.**
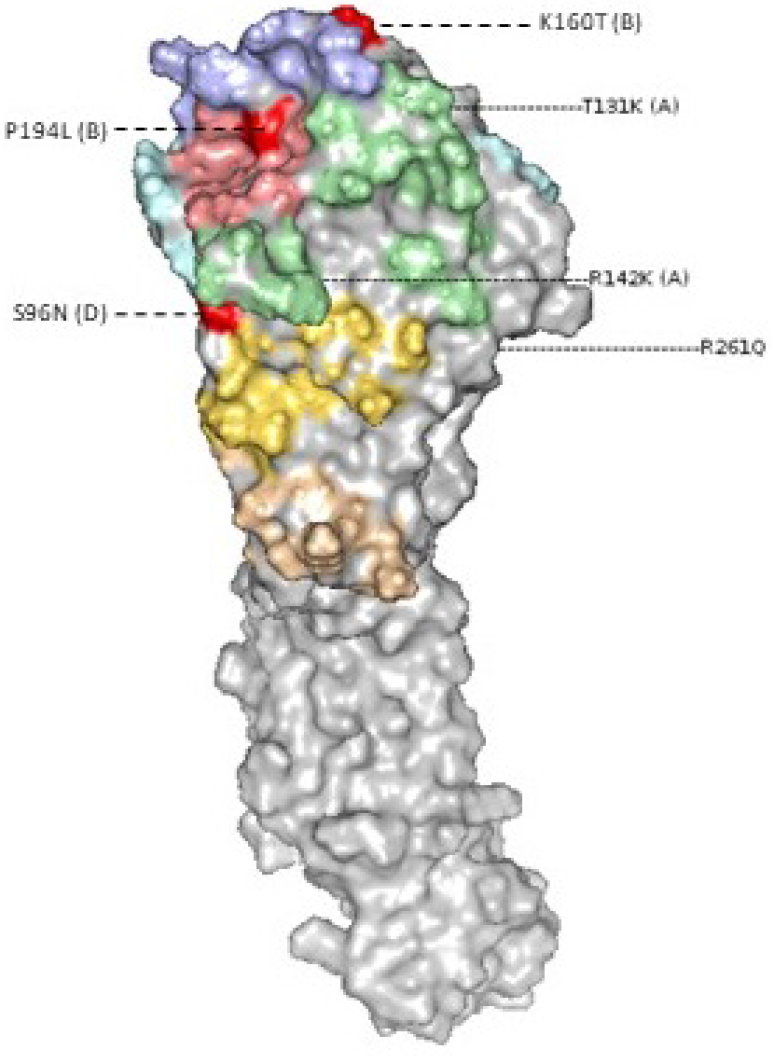
Differences in the hemagglutinin protein comparing A(H3N2) cell-propagated A/Washington/16/2017 to egg-propagated A/Hong Kong/4801/2014. Amino acid changes highlighted in red on the HA structure are those associated with egg-adaptation in A/Hong Kong/4801/2014. Antigenic sites are indicated in parenthesis.

### Serologic studies in influenza A(H3N2) cases and controls

Serologic analyses were performed on specimens collected from the subset of participants enrolled between November 27, 2017 and January 31, 2018, including all 15 influenza A(H3N2) infected cases identified by that date and 15 uninfected controls. Participants ranged in age from 20 to 93 years (median: 37 years), 53% were female, 63% were white race, 60% received an influenza vaccine, 38% reported fair or poor general health, and 41% were frail. Cases and controls were comparable in terms of age, sex, race, and date of enrollment (Table 2). Control subjects were more likely to report fair or poor general health, were more likely to be frail, and were more likely to be vaccinated; however, these differences were not statistically significant. Serum specimens were collected<5 days from illness onset from 73% of cases and controls.

**Table 2.** Characteristics of hospitalized influenza A(H3N2) infected cases and uninfected controls.

| Characteristic | Influenza A(H3N2) Case | Control | *P*-value |
| --- | --- | --- | --- |
| Age (20-93 years) – median (IQR) | 34 (22, 63) | 37 (28, 62) | 0.62 |
| Sex – no. (%) |  |  | 0.46 |
| Female | 9 (60) | 7 (47) |  |
| Male | 6 (40) | 8 (53) |  |
| Race/ethnicity – no. (%) |  |  | 0.70 |
| White (not Hispanic) | 10 (67) | 9 (60) |  |
| Other | 5 (33) | 6 (40) |  |
| Reported general health status – no. (%) |  |  | 0.32 |
| Excellent/very good/good | 10 (71) | 8 (53) |  |
| Fair/poor | 4 (29) | 7 (47) |  |
| Frailty status – no. (%) |  |  | 0.18 |
| Not frail | 10 (71) | 7 (47) |  |
| Frail | 4 (29) | 8 (53) |  |
| Illness onset to blood collection – no. (%) |  |  | 1.00 |
| 0-4 days | 11 (73) | 11 (73) |  |
| 5-9 days | 4 (27) | 4 (27) |  |
| Influenza vaccination status – no. (%) |  |  | 0.14 |
| Vaccinated | 7 (47) | 11 (73) |  |
| Unvaccinated | 8 (53) | 4 (27) |  |
| Date of Enrollment – median (IQR) | 1/18/18 (1/8/18, 1/27/18) | 1/9/18 (12/19/17, 1/22/18) | 0.12 |

Both neutralizing and NAI antibody titers to various influenza antigens in combined cases and controls were first determined. Overall titers measured by MN were highest against the egg-grown A/HongKong/4801/2014 3C.2a vaccine strain (GMT [95% CI]: 97 [53, 177]). Titers against cell-grown strains were lower than against the egg-grown strain, and titers were comparable between cell-grown A/HongKong/4801/2014 vaccine strain and the four influenza A(H3N2) strains representative of those that circulated: A/Washington/16/2017 3C.2a2, A/Singapore/Infimh-16-0019/2016 3C.2a1, A/Maryland/53/2017 3C.2a1b+135N, and A/Wisconsin/327/2017 3C.2a1b+135K (GMT range: 19-34). NAI GMTs were >40 against both A/HongKong/4801/2014 (GMT [95% CI]: 54 [33, 89]) and A/Singapore/Infimh-16-0019/2016 (GMT [95% CI]: 41 [24,69]).

The correlation of titers against egg- and cell-grown viruses in combined cases and controls was next evaluated. Egg-grown A/HongKong/4801/2014 titers were most strongly correlated with cell-grown A/HongKong/4801/2014 (ρ=0.71) and A/Wisconsin/327/2017 titers (ρ=0.70) and less correlated with A/Singapore/Infimh-16-0019/2016 (ρ=0.56) and A/Washington/16/2017 titers (ρ=0.53) (Supplemental Figure 1). Correlations between antibody titers against each of the cell-grown viruses were higher (ρ range: 0.80–0.90) with each other than against the egg-grown A/HongKong/4801/2014 virus.

### Relation of antibody titers to vaccination status

There were 18 individuals who had been vaccinated, and the specific influenza vaccine they received could be confirmed for 15. All 15 individuals with confirmed vaccine product information received vaccines containing inactivated egg-grown virus; 10 received quadrivalent standard dose vaccines (9 GSK, 1 Seqirus), 3 received trivalent high dose vaccines (Sanofi Pasteur), 1 received trivalent adjuvanted vaccine (Seqirus), and 1 received trivalent unadjuvanted vaccine (Seqirus). Vaccinated individuals had significantly higher titers against the egg-grown A/HongKong/4801/2014 vaccine strain than unvaccinated individuals (GMT: 173 vs 41; *P* = 0.01). For all other viruses, including the cell-grown A/HongKong/4801/2014 vaccine strain, MN titers were similar comparing unvaccinated and vaccinated individuals (Figure 3). Interestingly, NAI titers were significantly higher for vaccinated individuals than they were for unvaccinated individuals (Figure 4) for both the A/HongKong/4801/2014 (GMT: 80 vs 30; *P* = 0.05) and A/Singapore/Infimh-16-0019/2016 (GMT: 63 vs 21; *P* = 0.03).

**Figure 3.**
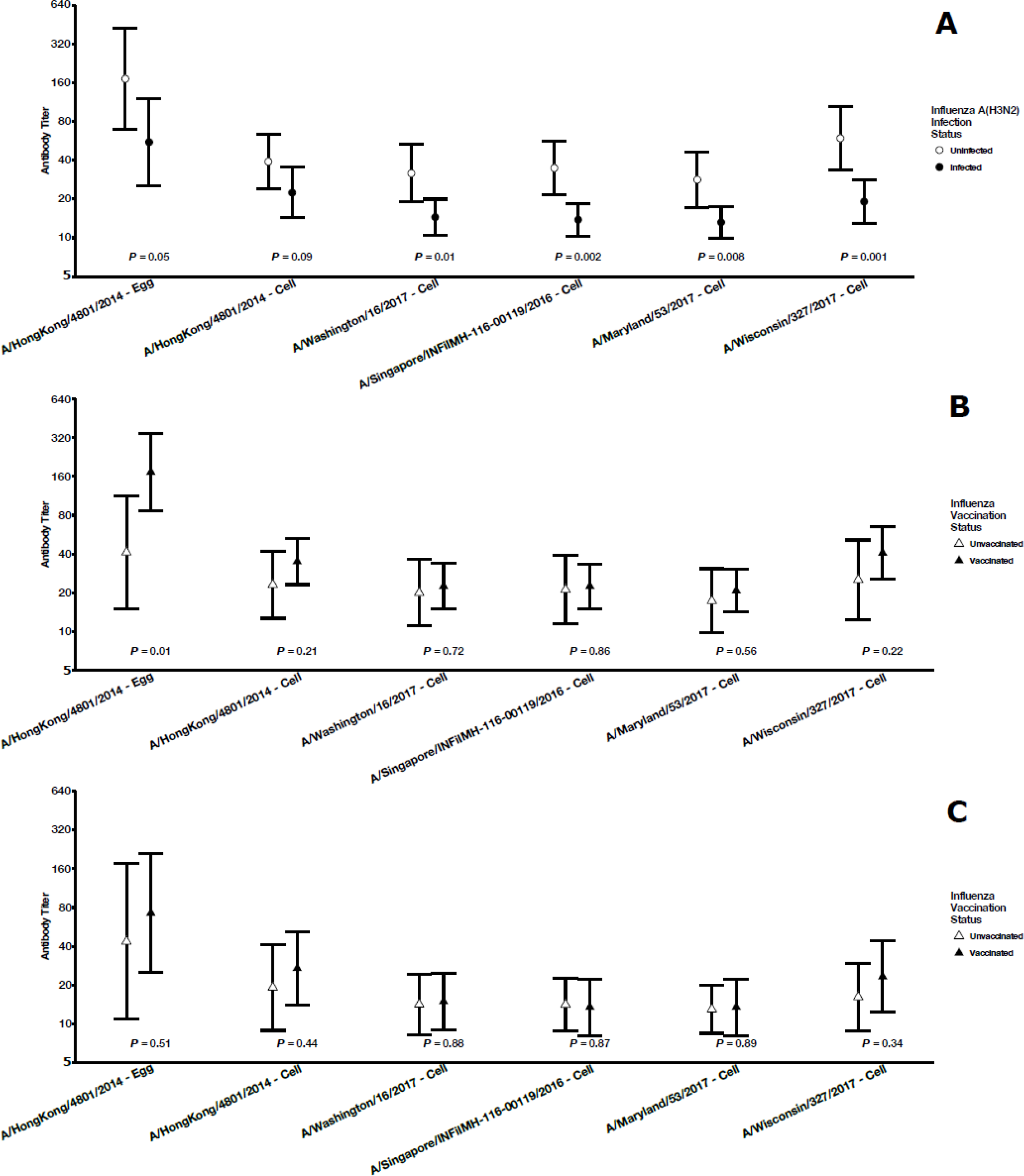
Geometric mean microneutralization titers by A) influenza A(H3N2) infection status, B) influenza vaccination status, and C) influenza vaccination status among influenza A(H3N2) infected cases.

**Figure 4.**
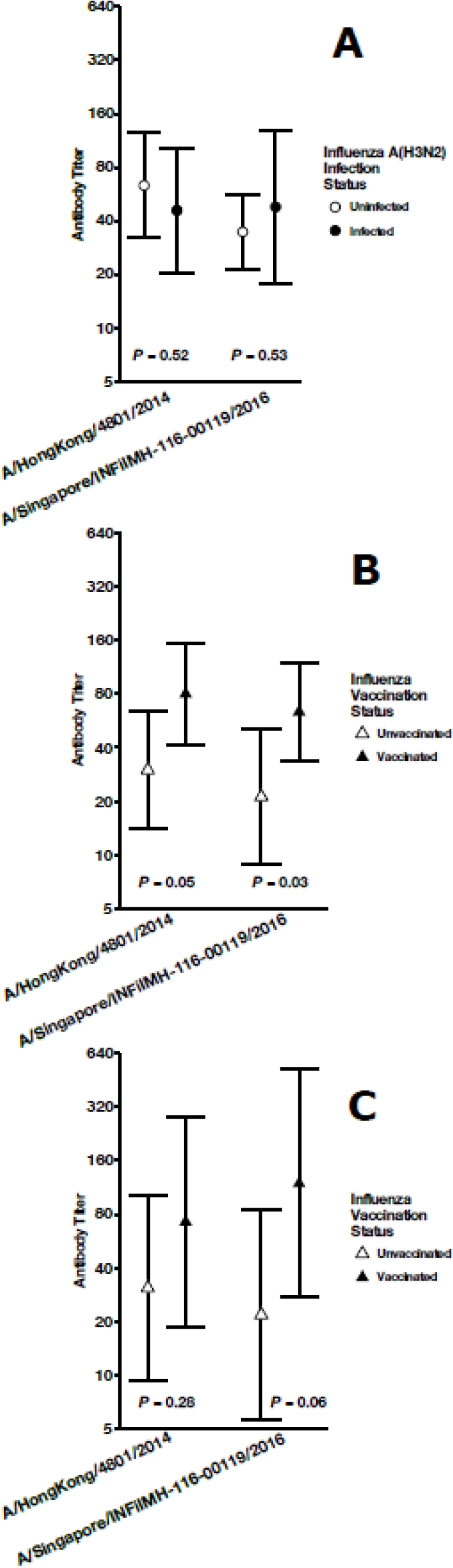
Geometric mean neuraminidase inhibition titers by A) influenza A(H3N2) infection status, B) influenza vaccination status, and C) influenza vaccination status among influenza A(H3N2) infected cases.

### Relation of antibody titers to infection status

Eight of the 15 individuals (53%) with influenza infection had been vaccinated against influenza in the 2017-2018 season. The majority of these vaccine failures had low MN titers against A/Washington/16/2017, the virus most similar to those 2a2 viruses that predominated in the region. For all target strains, uninfected controls had higher MN titers than influenza A(H3N2) infected cases (Figure 3). These differences between cases and controls were statistically significant for all cell-grown viruses representative of circulating viruses (all *P* < 0.01), but the differences were only marginally significant for antibody titers against egg- and cell-grown A/HongKong/4801/2014 vaccine strain viruses (respectively: *P* = 0.05 and *P* = 0.09). In contrast, NAI titers were not substantially different comparing cases and controls for either NA target (Figure 4).

We used multivariable logistic regression models to determine if the apparent association between antibody against egg-grown A/HongKong/4801/2014 and reduced odds of influenza A(H3N2) infection was independent of the reduction in infection associated with antibody against A/Washington/16/2017, the virus most similar to those that predominated in this population (Supplementary Table 2). In unadjusted models, MN titers against egg-grown A/HongKong/4801/2014 (odds ratio: 0.71; 95% CI: 0.50, 1.01) and A/Washington/16/2017 (odds ratio: 0.40; 95% CI: 0.19, 0.86) were both associated with reduced odds of infection, but only the effect for A/Washington/16/2017 was statistically significant. These effects correspond to an estimated 50% reduction in odds of infection for an individual with a titer of 20.3 against egg-grown A/HongKong/4801/2014 and 8.5 against A/Washington/16/2017 relative to an individual with undetectable titer for each virus, respectively (Figure 5A and B). The effects of both MN titers were somewhat attenuated and no longer statistically significant as estimated in adjusted models. After adjustment for titers against A/Washington/16/2017, the titer required for a 50% reduction in odds of infection increased to 95.3 for egg-grown A/HongKong/4801/2014 (Figure 5C). After adjustment for titers against egg-grown A/HongKong/4801/2014, the titer required for a 50% reduction in odds of infection increased to 9.3 for A/Washington/16/2017 (Figure 5D).

**Figure 5.**
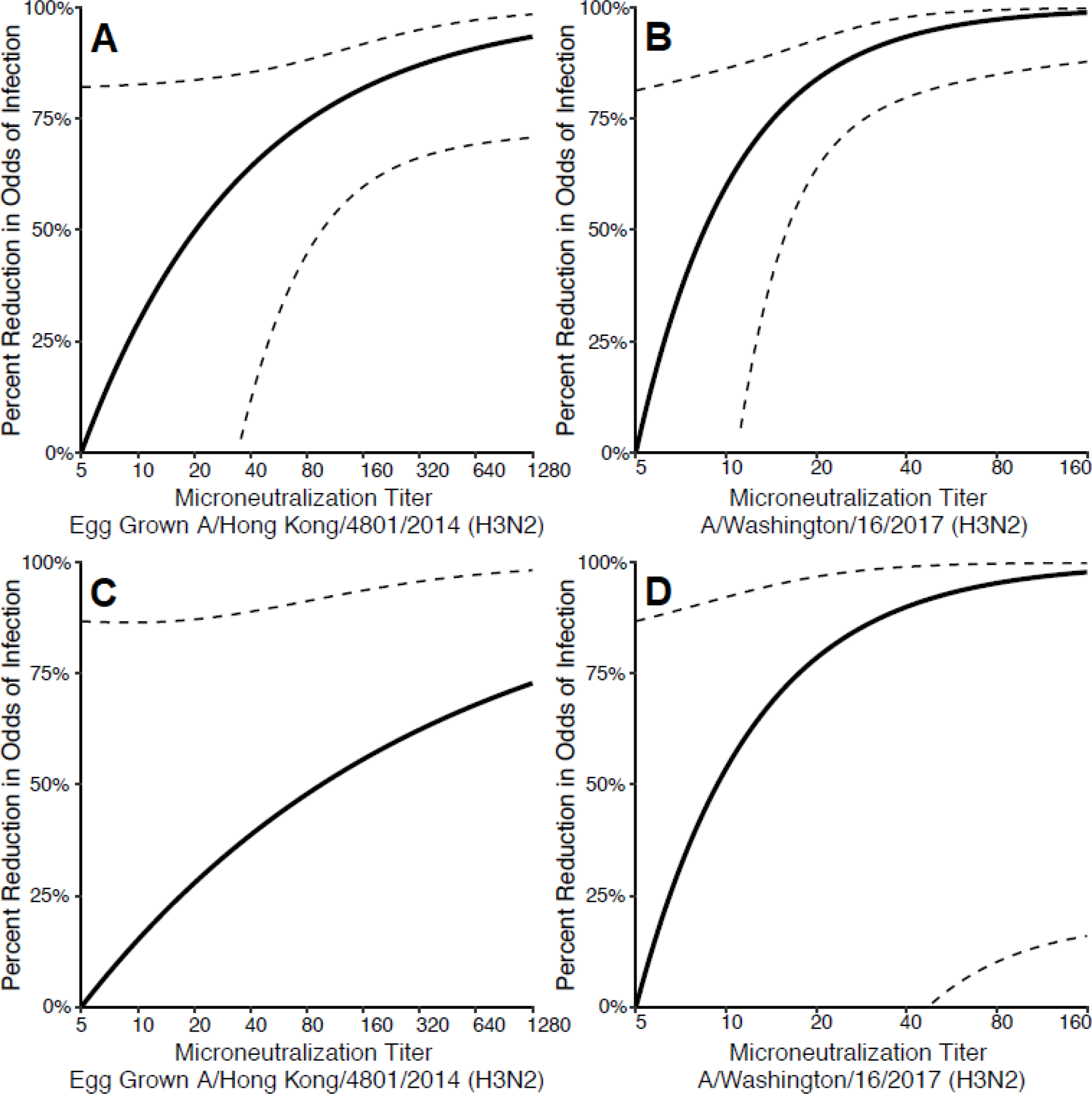
Estimated percent reduction in odds of influenza infection by increasing microneutralization titers against egg-propagated influenza A/Hong Kong/4801/2014 (H3N2) and A/Washington/16/2017 (H3N2). Unadjusted estimates for each virus target are presented in panel A and B; the estimates presented in panels C and D adjust for titer against the other virus. 95% confidence intervals indicated by dotted lines.

## DISCUSSION

As in other parts of the United States, the intense local influenza A(H3N2) season was predominately 3C.2a2[21]. These and other 3C.2a descendant viruses were well inhibited by ferret-antisera raised against representative 3C.2a viruses grown in cell culture, but were not as well inhibited by antisera raised against egg-grown viruses[9]. There was some circulation of 3C.3a viruses, but not during the main part of the outbreak. Therefore, we focused our analysis on those 3C.2a viruses that were representative of those included in the vaccine and that predominantly circulated. This allowed us to focus on the role of egg adaptation, which had been hypothesized in laboratory studies to be responsible for the reduced VE in 2017-18 [6,7]. We were able to provide evidence that the antibody response to influenza vaccination was specific to egg-adapted influenza A(H3N2) strains and that these antibodies did not strongly correlate with protection from infection during the 2017-2018 season.

In this hospitalized population, MN titers against the egg-adapted influenza A/HongKong/4801/2014 (H3N2) strain included in the 2017-2018 influenza vaccine were significantly higher in vaccinated compared to unvaccinated individuals. However, both vaccinated and unvaccinated groups had similar titers against cell-grown A/HongKong/4801/2014 (H3N2). This suggests that the antibody responses to vaccination in this population are highly specific to the egg-adapted strain with very little cross-reactivity to the circulating virus. Although not statistically significant, there was an indication that these antibodies against egg-adapted influenza A/HongKong/4801/2014 (H3N2) strain provided some level of protection against infection. This may be consistent with low, but non-zero, estimated effectiveness of the 2017-2018 influenza vaccine. In contrast, titers against wild-type influenza A(H3N2) 3C.2a1 and 3C.2a2 viruses were lower overall in this population, but they were significantly correlated with protection against infection during the 2017-2018 season. These results were consistent even in models accounting for correlation between antibodies against specific strains.

Immunologic studies have proposed mechanisms for the lack of antibody cross-reactivity between egg-grown and circulating viruses. A study of plasmablasts in individuals receiving influenza A(H1N1)pdm2009 vaccine found that individuals with a highly-specific antibody response to a reversion present in the vaccine strain had no binding to the A(H1N1) circulating strain[22]. With regards to more recent H3N2 strains, similar immunologic patterns of antibody misdirection have been reported for residue 160 in antigenic site B of HA[7]. Influenza A(H3N2) 3C.2a viruses that have circulated since the 2014-2015 season possess a K160T mutation which introduces a glycosylation to antigenic site B that was not present prior to 2014-2015. This mutation is reversed in the corresponding egg-propagated 3C.2a egg-adapted strain present in the 2016-2017 and 2017-2018 vaccines, thus allowing the immune system to focus the antibody response on this epitope which is otherwise shielded in circulating strains[7].

Antibody against the neuraminidase of A/HongKong/4801/2014 and A/Singapore/INFIMH-16-0019/2016 was not associated with protection against infection. This is in contrast to past studies, which found that antibody to the neuraminidase can play a role in protection, including during the 1968 pandemic [23–25]. A lack of association between anti-neuraminidase antibody-mediated immunity and protection could explain, in part, the high incidence of infection during the 2017-2018 influenza season. We have previously demonstrated that anti-neuraminidase antibody was not associated with protection during the 2014-2015 season, which was also characterized by high incidence of influenza A(H3N2) infections [26]. Although neuraminidase content is not standardized in influenza vaccines, we observed that anti-neuraminidase antibody titers were higher among vaccinated individuals suggesting that some of the existing anti-neuraminidase antibody in this population resulted from vaccination. This is consistent with previous studies that have directly observed a neuraminidase directed antibody response following vaccination [23,27]. Given the lack of protection demonstrated and questions about the quality of vaccine induced neuraminidase antibody [28], it is clear that more work needs to be done to understand antigenic drift of the neuraminidase, and how and when anti-neuraminidase antibodies mediate protection.

There are now at least two influenza vaccines available in the United States that are not produced in eggs. However, evidence for the effectiveness of these vaccines relative to those produced in eggs is limited given that few head-to-head trials have been performed. One of these trials was carried out during the 2014-2015 season with predominant circulation of antigenically drifted influenza A(H3N2), and found 30% higher relative efficacy in older adults for a recombinant hemagglutinin-based vaccine compared to standard egg-grown inactivated influenza vaccine [29]. Vaccines grown in eggs and in cell-culture were found to have similar efficacy overall in a placebo controlled trial carried out in 2007-2008, but only the cell-culture derived vaccine demonstrated statistically significant efficacy against influenza A(H3N2) [30]. Because these vaccines are still relatively new to the market and production remains limited, influenza vaccines produced in eggs represent the vast majority of those administered. This limited use currently precludes estimation of comparative effectiveness in large-scale observational studies.

The serum specimens in this study were collected at a single time point early during acute illness[26]. We have previously described evidence that antibody measured in acute specimens such as these reflects pre-infection immune status rather than a de novo response to infection. While we demonstrated differences in antibody titer by vaccination status, we are unable to measure the magnitude of antibody response to vaccination in the absence of a paired pre-vaccination specimen. In addition, this analysis included a relatively small number of specimens from a single hospital. Because of the severity of the 2017-2018 epidemic was evident early in the season we prioritized rapid serologic testing of the specimens that were available to date during first half of the season. These two factors may limit the broad generalizability of the results; however, the influenza viruses that circulated locally overall and during the analysis period were similar to those that predominated nationwide during the same season. The small sample size also precluded our ability to perform additional sub-analyses to assess whether results might vary within specific age groups or by history of prior vaccination.

In one sense, the 2017-2018 influenza vaccine appears to have worked as intended: vaccinated individuals had high antibody titers against the influenza strain included in the vaccine. While this may have provided some protection, it is clear that the antibody response to vaccination was not specific enough to the viruses that actually circulated to provide high levels of protection. This was likely the result of changes to the vaccine strain virus from egg-adaptation. Available influenza vaccines that are not manufactured in eggs may provide some hope of increased effectiveness in future years. Unfortunately, these vaccines are still dependent on selecting viruses that are close antigenic matches to those that will circulate and are subject to the limitations of annual strain selection timelines. This underscores the need for the development of the next-generation of influenza vaccines that provide better, broader, and longer lasting protection [31].

---

**Supplementary Table 1.**
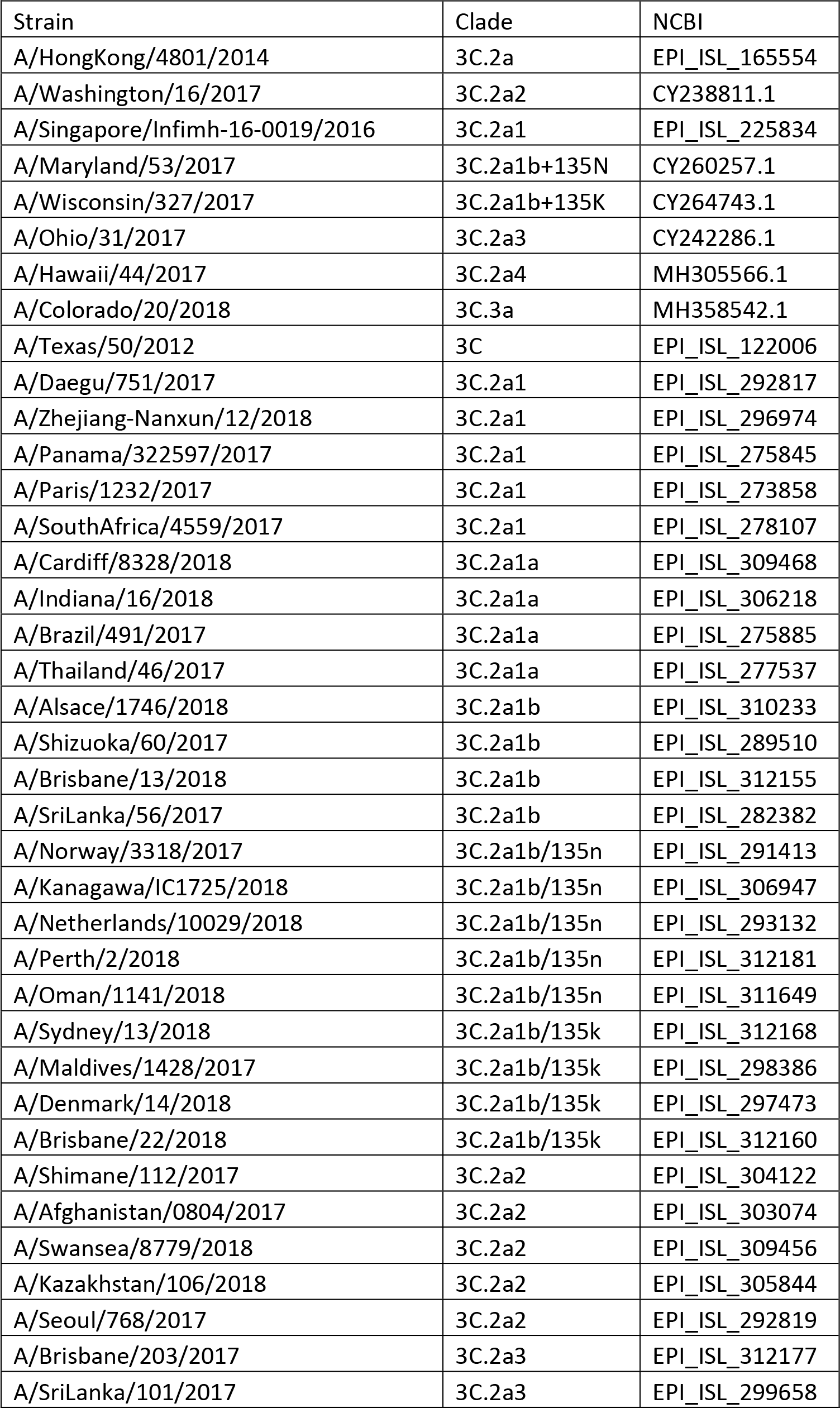
Accession numbers of reference strains and study sequences used in phylogenetic analysis.

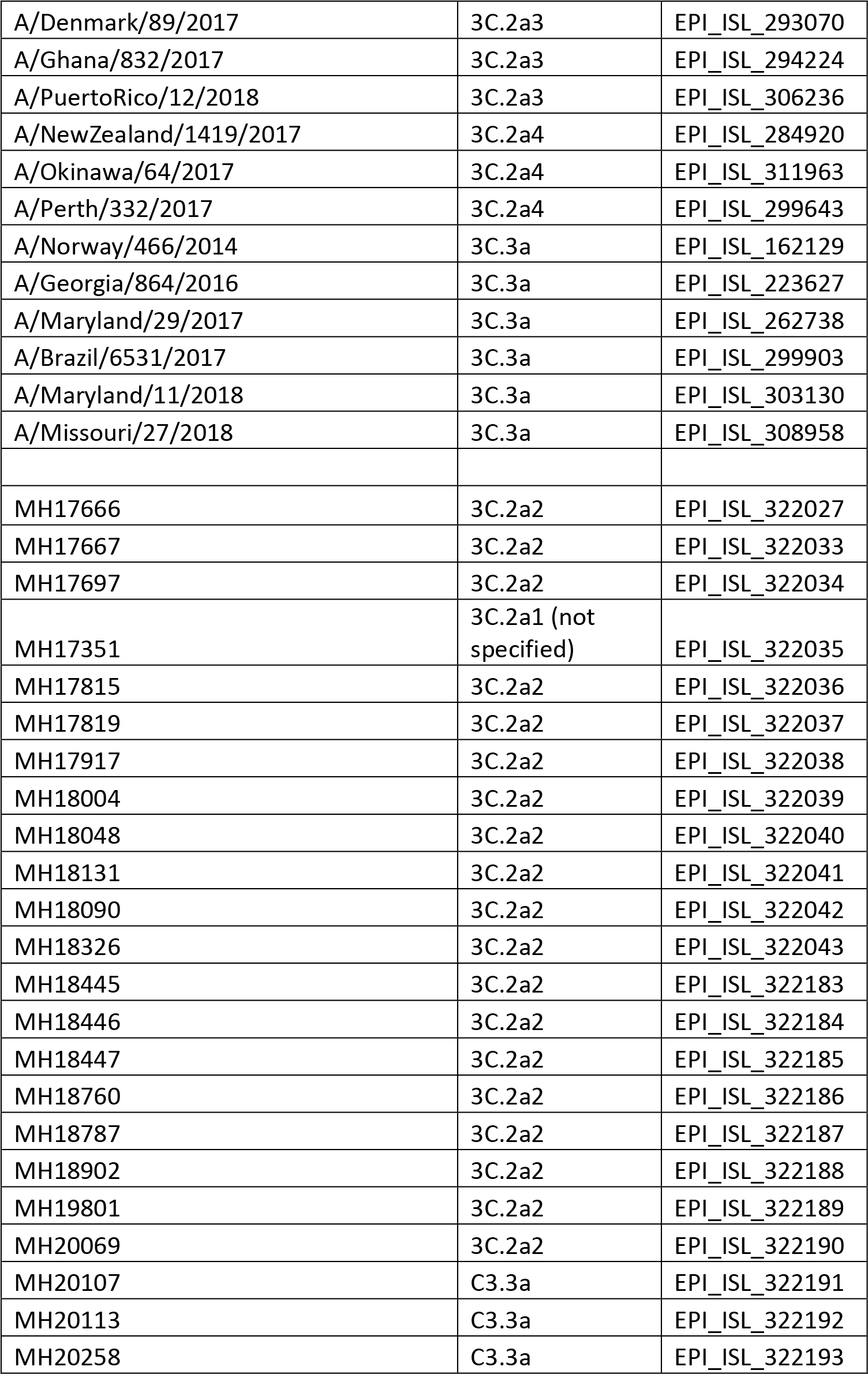

**Supplementary Table 2.**
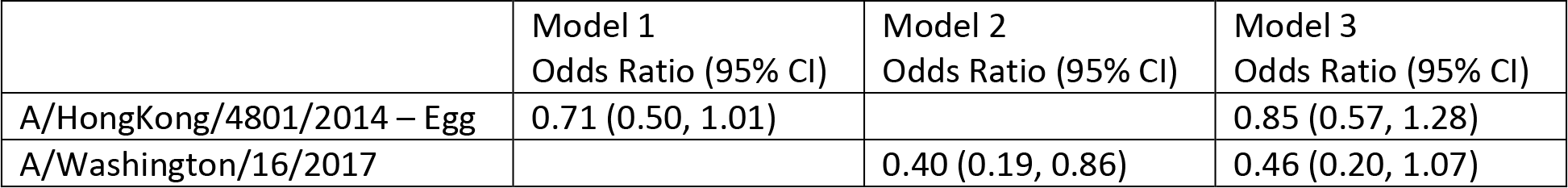
Estimated reduction in odds of infection associated with a 2-fold increase in microneutralization titers against egg-propagated influenza A/Hong Kong/4801/2014 (H3N2) and A/Washington/16/2017 (H3N2). Unadjusted estimates for each virus target were estimated in models 1 and 2; model 3 estimated the reduction in odds of infection associated with each strain-specific titer adjusted for the other.

**Supplementary Figure 1.**
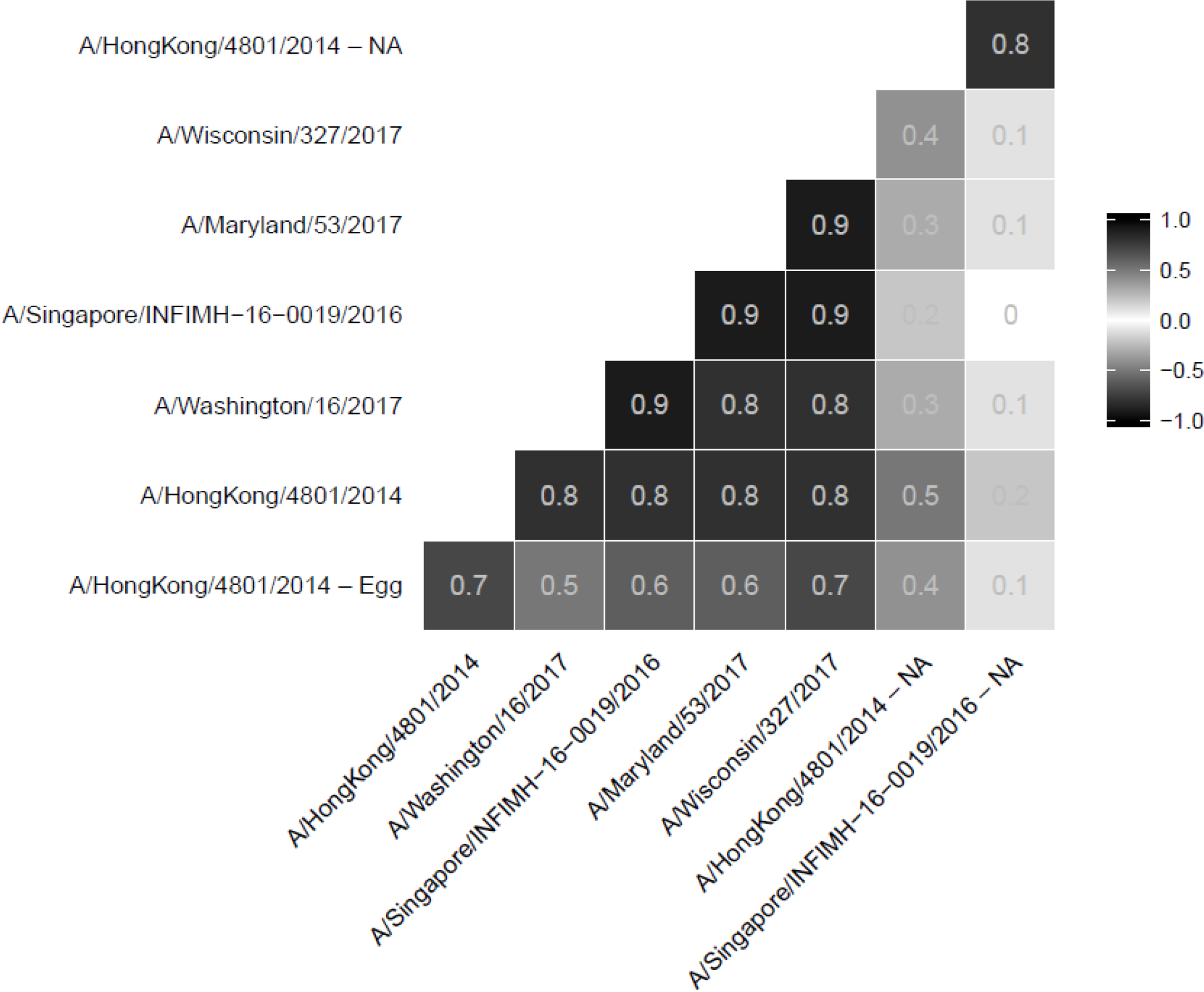
Correlation between microneutralization and neuraminidase inhibition antibody titers against egg- and cell-propagated influenza A(H3N2) viruses.

## Acknowledgements

We thank Maryna Eichelberger, Hongquan Wan, Jin Gao, and Laura Couzens (Food and Drug Administration) for technical support and providing reassortant influenza viruses for use in the enzyme-linked lectin assays. St Jude Children’s Research Hospital provided plasmids that were used to generate these reassortant influenza viruses. We thank Mrs F Liaini Gross, Lauren Horner and Makeda Kay from Influenza Division, Centers for Disease Control and Prevention for technical support for virus propagation and specimen management.

